# Rapid sex-specific evolution of age at maturity is shaped by genetic architecture in Atlantic salmon

**DOI:** 10.1101/317255

**Authors:** Y. Czorlich, T. Aykanat, J. Erkinaro, P. Orell, CR. Primmer

## Abstract

Understanding the mechanisms by which populations adapt to their environments is a fundamental aim in biology. However, it remains challenging to identify the genetic basis of traits, provide evidence of genetic changes and quantify phenotypic responses. Age at maturity in Atlantic salmon represents an ideal trait to study contemporary adaptive evolution as it has been associated with a single locus in the *vgll3* region, and has also strongly changed in recent decades. Here, we provide an empirical example of contemporary adaptive evolution of a large effect locus driving contrasting sex-specific evolutionary responses at the phenotypic level. We identified an 18% decrease in the *vgll3* allele associated with late maturity (*L*) in a large and diverse salmon population over 36 years, induced by sex-specific selection during the sea migration. Those genetic changes resulted in a significant evolutionary response in males only, due to sex-specific dominance patterns and *vgll3* allelic effects. The *vgll3* allelic and dominance effects differed greatly in a second population and were likely to generate different selection and evolutionary patterns. Our study highlights the importance of knowledge of genetic architecture to better understand fitness trait evolution and phenotypic diversity. It also emphasizes the potential role of adaptive evolution in the trend toward earlier maturation observed in numerous Atlantic salmon populations worldwide.

## Introduction

Understanding the mechanisms by which populations adapt to their environments is a fundamental aim in biology^1,2^. Such mechanisms may represent the only way for certain populations to persist in the face of strong human pressures and accelerated rates of climate change altering their environment. Temporal monitoring has documented recent and rapid phenotypic changes in wild populations in many species ^e.g. 3,4^ However, whether or not such phenotypic changes are adaptive often remains unclear ^5,6^. Obtaining evidence of adaptive evolution requires knowledge of the genetic basis of traits and subsequent demonstration that natural selection induces changes in this genetic component ^6^. Although the ideal strategy for demonstrating adaptive evolution is to study the genes directly controlling the traits under selection, such examples are extremely scarce ^6,7^. Despite the increased availability of genomic data, the identification of large-effect loci controlling phenotypes of ecological significance and understanding how contemporary selection affects the allele frequency of such genes remains indeed challenging^8^. In cases where the genetic architecture of a trait is well characterized e.g. when a large-effect locus has been identified, retrospective genetic analyses of archived material for the gene(s) controlling the trait in question can be performed and provide detailed information about its evolutionary dynamics ^e.g^. ^9^.

Age at maturity in Atlantic salmon, defined here as the number of years spent at sea prior to maturation, has recently been shown to be associated with a single large-effect locus with sex-specific effects, located within a narrow (<100kb) region around the *vgll3* gene ^10^. The same locus has also recently been linked with gender-biased auto-immune diseases in humans 11. In Atlantic salmon, age at maturity reflects a classic evolutionary trade-off, as larger, later-maturing individuals typically have higher reproductive success, but run a greater risk of mortality before first reproduction. Sex-specific selection optima may exist for this trait ^10^. Males generally mature earlier and at smaller size, whereas females mature later and have a stronger correlation between body size and reproductive success compared with males^12^. It was suggested that the sex-dependent dominance observed at *vgll3* partially resolves this sexual conflict ^10,13^. Furthermore, the age structure of many salmon populations has changed worldwide in recent decades, generally towards an increasing proportion of smaller, earlier maturing individuals ^e.g. 14,15 but see 16^. However, the reasons for this, and whether it is an adaptive change, remain unknown^17^. Therefore, age at maturity in Atlantic salmon provides a rare opportunity to investigate the contemporary change of a phenotypic trait directly at the genetic level.

We studied a 40 year time series of two closely related Atlantic salmon populations from northern Europe with contrasting maturation age structure. Despite a low level of genetic divergence between them (*F*_ST_ = 0.012)^18^, one population (Tenojoki) displays a high level of life-history diversity including a high proportion of large, later maturing individuals in both sexes, whereas the other population (Inarijoki) consists primarily of individuals of younger maturation ages, and with less life-history variation, particularly in males ^15^. Here, we utilized a 40 year time series to detect potential signs of adaptive evolution in age at maturity by contrasting allele frequency changes at the maturation-linked gene *vgll3* with life-history phenotypes in 2500 samples from the two populations. We also investigated the occurrence of sex- and population-specific genetic architecture and selection, potentially explaining the observed diversity variation in age at maturity.

## Results

### Temporal changes in age at maturity

We first quantified temporal phenotypic changes in both populations. There was a non-linear decrease in the age at maturity of Tenojoki individuals, with the mean maturation age of males declining by >40% (from 2.2 to 1.3 years; *edf* = 3.87, *F* = 5.11, P < 0.001) and of females by 8.1% (from 3.0 to 2.7 years; *edf* = 1.27, *F* = 0.57, P = 0.02), during the 36 year time period (Figure 1**a**). In Tenojoki males, the decrease occurred primarily between 1971 and 1987 before stabilization, while in females, age at maturity gradually decreased over the 36 year study period, explained best by a slightly nonlinear slope (Figure 1**a**). In comparison, Inarijoki males were virtually devoid of variation in age at maturity, with almost all males having spent one year at sea before maturing (*edf* = 0.00, *F*=0.00, P = 0.731), whereas mean age at maturity in females fluctuated cyclically over the 37 years (*edf* = 10.14, *F* = 6.035, P < 0.001, Figure 1**b**), but with no indication of a decrease in average maturation age (Figure 1**b**).

**Figure 1:**
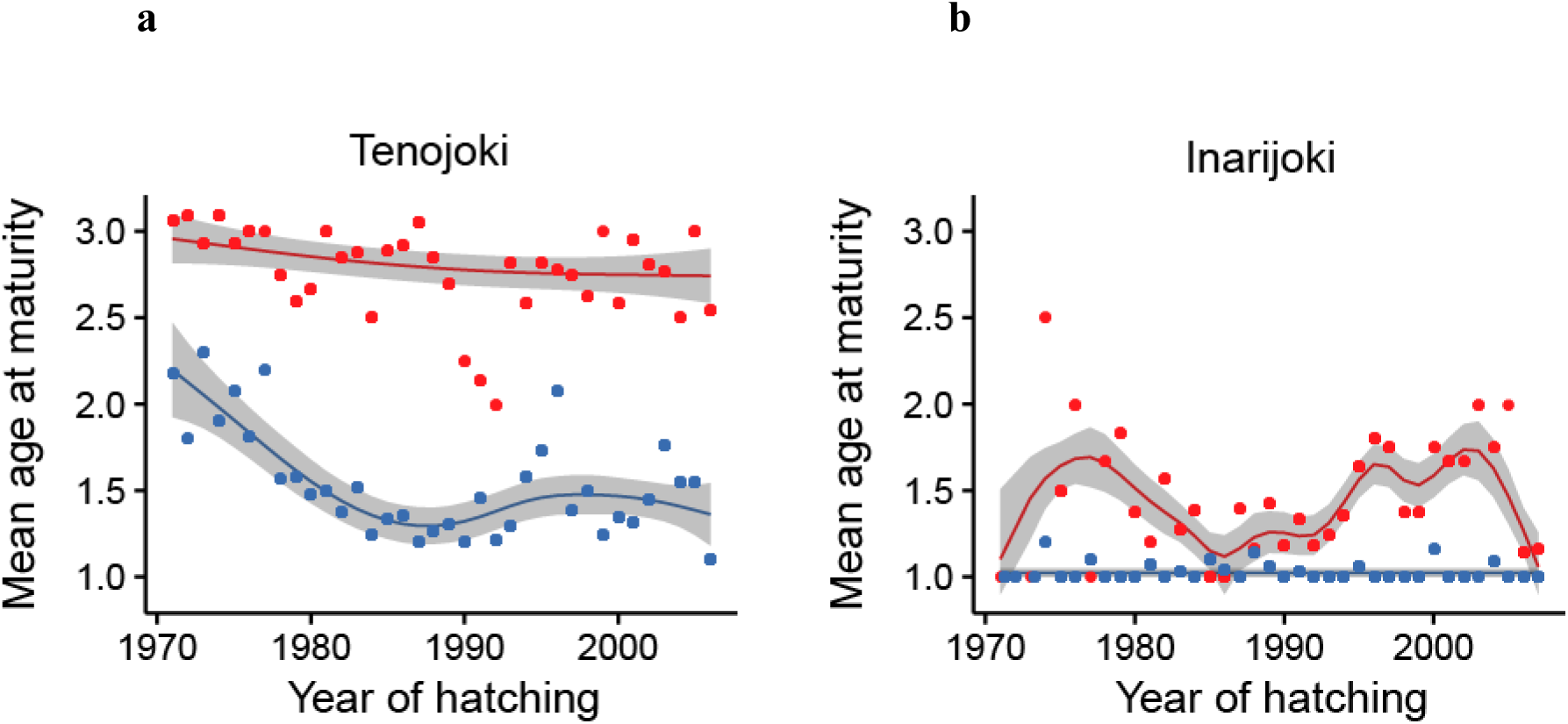
Change in mean age at maturity in the a, Tenojoki and b, Inarijoki populations. Females are in red (*N* Tenojoki = 467, *N* Inarijoki = 261) and males in blue (*N* Tenojoki = 699, *N* Inarijoki = 570). Lines represent fitted values from the GAM ± 1.96 SE, points are observed annual means.

### Genetic architecture of age at maturity

We hereafter use genetic architecture to refer to the additive and dominance effects of *vgll3* on age at maturity. The *vgll3* genotypes had a sex-specific effect on the probability to observe the different ages at maturity in the Tenojoki population (*χ*^2^_(6)_ = 27.58, P < 0.001). A sex-specific dominance pattern was observed in this population; heterozygote males had a mean age at maturity closer to homozygote *EE* (estimated dominance *δ_M_* = 0.09, CI_95_ = [0.02, 0.17], see Method) whereas heterozygote females had a phenotype closer to homozygotes *LL* (estimated dominance *δ_F_* = 0.80, CI_95_ = [0.65, 0.92]; Figure 2**a**). In the Inarijoki population, the *vgll3* genotypes were significantly associated with the probability to observe the different age at maturity groups (*χ*^2^_(4)_ = 56.41, P <0.001) but not in a sex-specific manner (*χ*^2^_(4)_ = 8.27, P = 0.08; estimated dominances of 0.13, CI_95_ = [0.05, 0.31] and 0.32, CI_95_ = [0.17, 0.50] in Inarijoki males and females, respectively). Differences in mean age at maturity between homozygotes varied depending on the sex and population (i.e. additive or allelic effect: effect of the substitution of one allele for the other). In the Tenojoki population, the relative difference in mean age at maturity between alternative *vgll3* homozygotes was about three times higher in males (+106% for *LL*, +1.17 years, CI_95_ = [0.99, 1.33]) than in females (+32% for *LL*, +0.71 years, CI_95_ = [0.51, 0.91]). This pattern was inverted in Inarijoki, with the relative difference in mean age at maturity between female homozygotes being about six times larger (+74% for *LL*, +0.94 years, CI_95_ = [0.68, 1.25]) than in males (+12% for *LL*, +0.13 years, CI_95_ = [0.05, 0.22], Figure 2**b**). These results imply that selection during the sea migration, defined as the relative difference in survival between genotypes, is likely to vary between sexes and populations. There was no statistically significant change in the effect size of *vgll3* on maturation age over time in either population (Tenojoki: *χ*^2^_(6)_ = 6.07, P = 0.42; Inarijoki: χ^2^_(4)_ = 4.41, P = 0.35).

**Figure 2:**
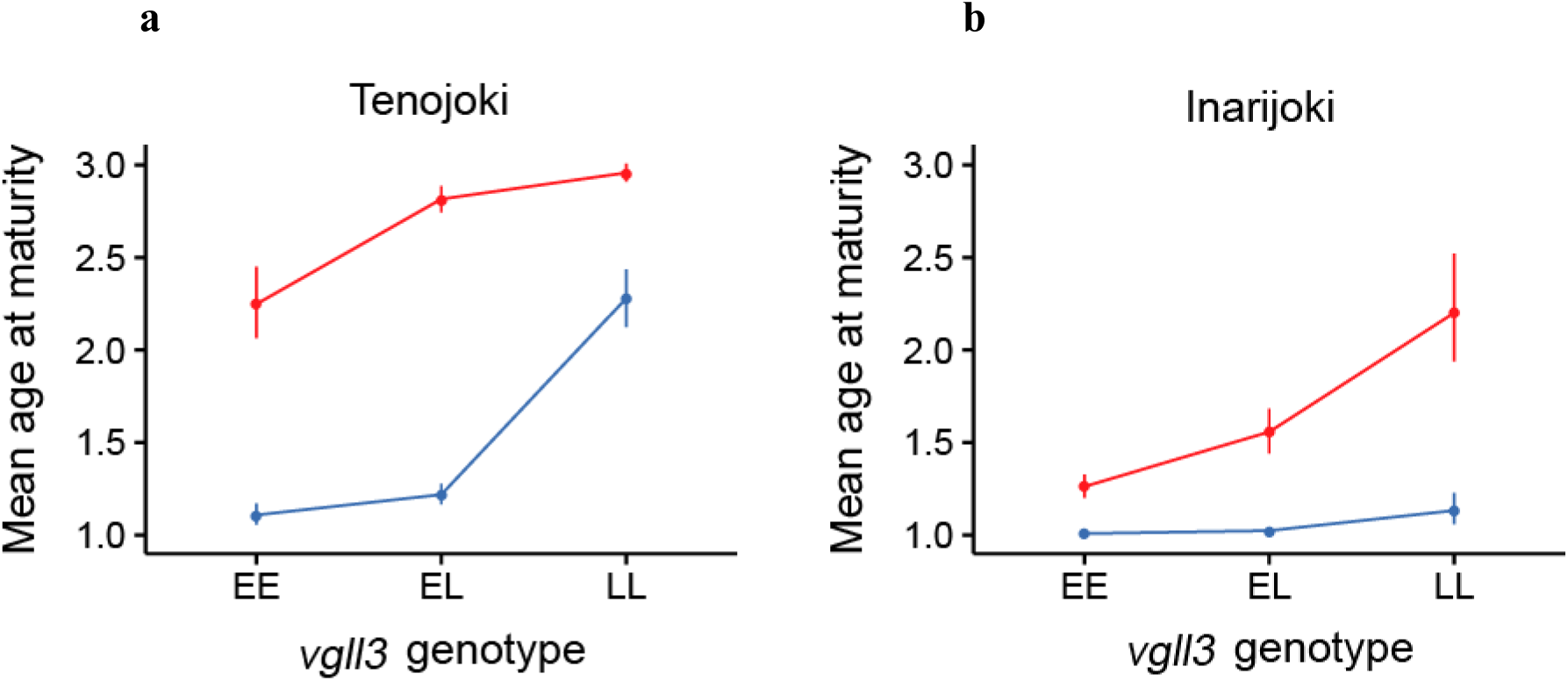
Mean age at maturity as a function of *vgll3* genotype in the a, Tenojoki and b, Inarijoki populations. Females are in red (*N* Tenojoki = 522, *N* Inarijoki = 286) and males in blue (*N* Tenojoki = 804, *N* Inarijoki = 612). Means are calculated from multinomial models fitted values, averaged over years. Error bars represents 95% bootstrap confidence intervals based on 1000 replicates.

### Evolution of *vgll3* and signals of selection

The *vgll3* late maturing (*L*) allele frequency decreased significantly, from 0.66 to 0.54 (18%) in 36 years, in the Tenojoki population (*F*(*_1_*) = 7.80, P = 0.009; log-odd slope = −0.014 (CI_95_ = [−0.004, −0.024]; Figure 3**a**). This allele frequency change was the highest of all the 144 genome-wide SNPs assessed and could not be explained by drift alone (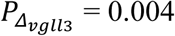, Figure 3**a**), nor after accounting for sampling variance (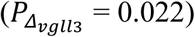. This observation provides strong support for natural selection acting against the *vgll3 L* allele in the Tenojoki population (see supplementary material). In the Inarijoki population, the trend in the *vgll3 L* allele frequency was also negative (log-odd slope ≈ −0.009, CI_95_ = [−0.023, 0.006] but not significant (*F*(_1_) = 1.29, P = 0.26, Figure 3**b**). About 10% of the 134 genome-wide SNPs assessed had a larger change in allele frequency than *vgll3* in this population (Figure 3**b**). Consequently, we could not rule out drift as the basis of this change (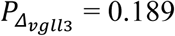 for drift, and 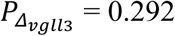 after accounting for sampling variance).

To further quantify the strength of selection driving changes in *vgll3* allele frequency, a Bayesian model was used to estimate selection coefficients whilst accounting for genetic drift, similar to a Wright-Fisher model (see Methods). The selection coefficient in the Tenojoki population was large and significantly higher than zero, albeit with large credibility intervals (−*s* = 0.33 (95% credibility interval = [0.01, 0.77], Supplementary Figure 1). In Inarijoki, there was no evidence for significant selection (*s* = 0.25, CI_95_ = [−0.08, 0.49]).

**Figure 3:**
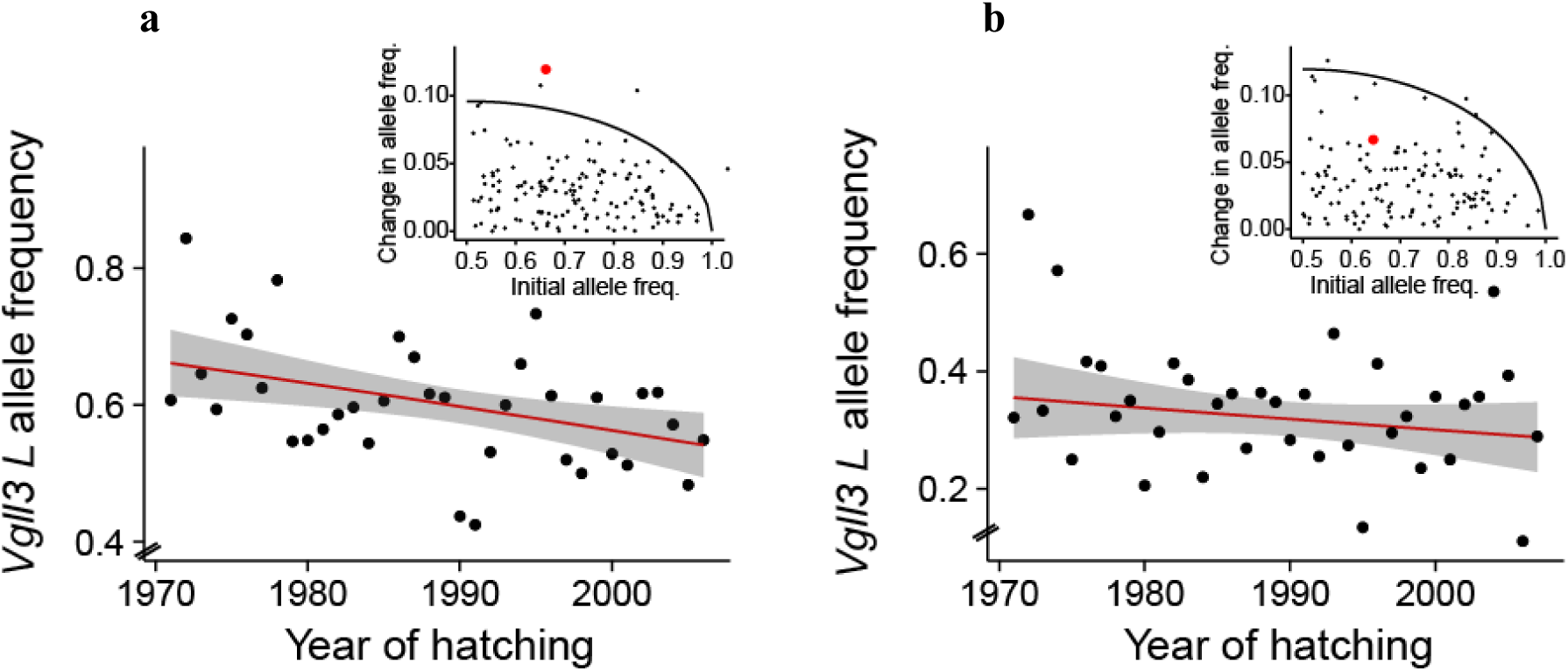
Temporal changes in *vgll3 L* allele frequency associated with late maturation in the a, Tenojoki and b, Inarijoki populations. The lines represent fitted values from the quasibinomial model with ± 1.96 SE (*N* Tenojoki = 1166, *N* Inarijoki = 765). The *vgll3* log-odd slope was estimated at −.014 (CI95 = [-0.004, −.024], P = 0.009) in Tenojoki and −0.009 (CI95 = [−0.023, 0.006], P = 0.26) in Inarijoki. Insets show the absolute estimated changes in allele frequencies of each SNP as a function of initial allele frequency in **a**, Tenojoki (144 loci) and **b**, Inarijoki (135 loci) over 36 and 37 years, respectively. The line represents the expected amount of drift at the 97.5 quantile. The *vgll3* locus is indicated in red.

The *vgll3 L* allele frequency differed between sexes in a contrasting manner in the two populations. The odds of possessing an *L* allele was 37% higher in females than in males in the Tenojoki population (CI_95_ = [0.12, 0.69], *F*_(1)_ = 8.72, P < 0.01, Figure 4) but 53% lower in Inarijoki (CI_95_ = [0.40, 0.65], *F*_(1)_ = 36.51, P < 0.001, Figure 4). This could be the result of either sex- and genotype-specific fertilization ratio or juvenile mortality in freshwater, or alternatively, selection at the *vgll3* locus differing between the sexes during sea migration, prior to returning to reproduce. In order to distinguish the latter possibility (selection at sea) from options involving selection during the freshwater phase we genotyped 143 and 108 juveniles of various ages collected from the same freshwater locations in Tenojoki and Inarijoki (1–3 years old, see Methods), respectively. Juvenile sex ratios were close to parity and the *vgll3 L* allele frequency was similar in both sexes in both populations (χ^2^_(1)_ = 3.27, P = 0.07 in Tenojoki, χ^2^_(1)_ = 0.04, P = 0.85 in Inarijoki, Figure 4). This provides support for the notion that selection strength acting on the *L* allele varies in a sex-specific manner during the marine life-history phase, as opposed to during the freshwater juvenile phase. Such sex-specific allele frequency patterns may be reinforced by sex-specific dominance (Supplementary Figure 2).

**Figure 4:**
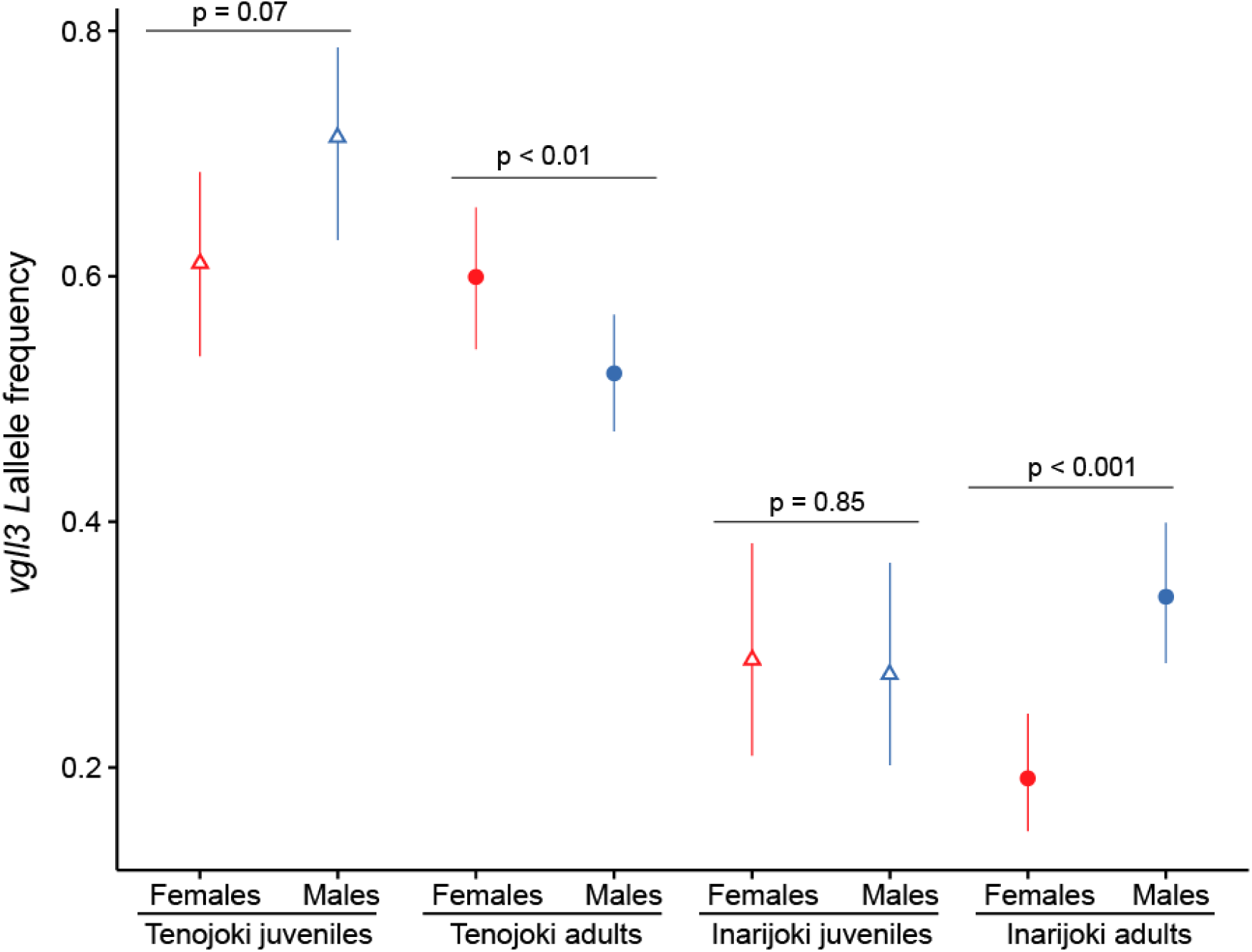
Model predicted mean *vgll3 L* allele frequency as a function of the sex and reproductive status in Tenojoki and Inarijoki. Error bars indicate 95% confidence intervals. Adult allele frequencies are from years 2006 and 2007 for females (red circles) and males (blue circles), respectively, in the Tenojoki and Inarijoki populations. Juveniles allele frequencies (triangles) are from 2012 in Tenojoki (2–3 years old) and 2016 in Inarijoki (1–3 years old).

### Sex-specific evolutionary response

In Tenojoki, the sex-specific genetic architecture drove contrasting evolutionary responses in the two sexes. Temporal changes in genotypes explained about 50% of the non-linear decrease in male age at maturity (0.46 years, *edf* = 3.95, *F* = 3.51, P < 0.001; Supplementary Figure 3) but didn’t explain the temporal changes in female age at maturity (0 years, *edf* = 0, *F* = 0.00, P = 0.54). Temporal changes in genotype frequencies were similar between sexes (χ^2^_(2)_ = 2.78, P = 0.25) and were thus unlikely to be the main driver of those sex-specific evolutionary responses. On the other hand, both the dominance patterns and *vgll3* effect sizes varied greatly between sexes (Figure 2**a**) and could contribute to the different responses. For instance, an individual with the *EE* genotype rather than *LL* matures, on average, 0.71 (CI_95_ = [0.51, 0.91]) or 1.17 (CI_95_ = [0.99, 1.33]) years earlier, depending whether it is a female or a male. The extent of the phenotypic response thus differs between the sexes for a similar *EE* genotype frequency change. Further, because of sex-specific dominance, the phenotypic response to a change in heterozygote frequency will also vary between the sexes. For example in Tenojoki females, the phenotypic change induced by a decrease in the *LL* genotype frequency would be partially compensated by the increase in the *EL* genotype frequency, which results in a similar phenotype distribution due to the dominance of the *L* allele (Figure 2**a**, Supplementary Figure 4). In contrast, in males, the recessivity of the *vgll3 L* allele (*δ_M_* = 0.09) would favor a larger decrease in age at maturity when the proportion of *EL* heterozygotes is increasing. Most of the observed decline in female age at maturity could be explained by the spawning year (0.20 year, *edf* = 2.16, *F*=21.61, P < 0.001). The year effect also explained a part of the decline in male age at maturity (0.15 year, *edf* = 2.126, *F* = 12.41, P < 0.001).

## Discussion

We provide convincing evidence of rapid adaptive evolution of age at maturity toward small, early-maturing individuals in a large Atlantic salmon population. This indicates that despite having a reproductive advantage due to their large size ^12,19^, the late maturation life-history strategy has become increasingly costly and modified the reproductive success vs survival trade-off such that earlier maturation is increasingly advantageous. Adaptive evolution may thus represent a realistic mechanism behind changes towards earlier age at maturity observed worldwide in the last decades in Atlantic salmon ^e.g^. ^14–16,20^ and other salmonid fish species ^e.g. 21^. What could be the causes of such rapid evolution of a life-history trait? One explanation is that it could be linked to recent rapid changes in the marine environment of the Teno salmon populations. For example, climate change may negatively affect Atlantic salmon marine growth and/or survival directly ^e.g. 22^ or indirectly through changes in Arctic food-webs and ecosystem functioning resulting from e.g. species range expansions^20,23,24^. Atlantic salmon occupying the northernmost parts of the globe will be unable to move to a colder climate in response to ocean warning, which would reinforce the importance of adaptation for population persistence. Another possibility is human-induced evolution of age at maturity through fishing targeting Atlantic salmon differentially according to their size, and therefore age at maturity ^e.g. 25,but see 26^, or reducing prey abundance ^e.g. 27^. Such environmental changes and/or human-induced pressure could negatively affect salmon survival at sea and thus increase the cost of late maturation, thereby potentially tipping the selective balance such that the size advantage at reproduction stemming from spending additional years at sea no longer compensates for the increased mortality and thus drives evolution towards younger maturation age. It is important to note however, that natural selection didn’t entirely explain the observed temporal changes in age at maturity in the Tenojoki population. Irrespective of the *vgll3* genotypes, the probability to mature at younger ages, after one or two years at sea, increased over time (Supplementary Information). This could be due to adaptive phenotypic plasticity^28^, through changes in maturation probability towards the same direction as selection, or to changes in allele frequencies of, as yet unknown, smaller effect loci. Further investigation is required to test these hypotheses. Regardless, such changes in population age structure can negatively affect the population growth rate and/or temporal stability induced via portfolio effects ^e.g. 29^ and also have negative consequences on genetic diversity levels ^e.g.30^ and thus are a concern for future population persistence.

Despite common temporal changes in *vgll3* allele frequency between the sexes, differing genetic architectures, in terms of additive and dominance patterns, contributed to sex-specific selection strengths and evolutionary responses to selection. We observed sex-specific differences in *vgll3* allele frequencies in adult salmon that were not present in pre-marine-migration juveniles from the same populations (Figure 4). Interestingly, the direction of the sex-specific differences was opposite in the two populations studied. The combined effects of sex-dependent dominance and sex-specific selection patterns can explain these contrasting patterns. Indeed, large between-populations variation in *vgll3* effects on age at maturity may influence selection and adaptive responses of individuals. The relative strength of allelic effects differed dramatically between sexes and these effects were in opposite directions in the two populations: in Inarijoki, the difference in mean age at maturity between homozygotes is about six times larger in females compared to males whereas in Tenojoki, the relative difference was three times higher in males (Figure 2). Therefore, selection against *LL* genotype individuals acts primarily on females in Inarijoki, but on males in Tenojoki (Figure 4, Supplementary material). However, sex-specific dominance also plays a role by introducing differences in allele frequencies between sexes that are dependent on population allele frequency (Supplementary Figure 2). Furthermore, sex-specific genetic architectures induce sex-specific evolutionary responses in Tenojoki, by accelerating the decrease in age at maturity in males and reducing the temporal phenotypic variation in females. Sex-specific dominance is likely to have evolved to reduce intra-locus sexual conflict ^10^. However, whether this genetic architecture is presently at its optimum is questionable in light of the quick decrease in *vgll3 L* allele frequency and age at maturity. Further studies are necessary to determine whether sexually antagonistic selection in Tenojoki is persisting in ever changing environments and to describe the extent, origin and consequences of among population variation in genetic architecture.

Age at maturity evolved rapidly under sex-specific selection in just 36 years, equivalent to 4–6 generations in Atlantic salmon. Despite being genetically similar, the two studied populations had distinctive genetic architectures, sex-specific selection and consequently *vgll3* allele frequencies variation. This study shows that variability in genetic architectures can create complex selection and evolutionary patterns between sexes and populations. This highlights the importance of determining the genetic basis of fitness traits in order to better understand their evolution and to explain the phenotypic diversity observed between populations and species.

## Material and methods

### Study site and sampling

The subartic Teno River forms the border between Finland and Norway and drains north into the Barents sea (68 - 70°N, 25–27°E). Genetically distinct salmon populations^31^ are distributed throughout the 16 386 km^2^ catchment area. Annual river catches range from about 20,000 to 60,000 individuals, representing up to 20% of the entire riverine Atlantic salmon harvest in Europe^32^. Atlantic salmon populations from Teno have been monitored since early 1970s with collections of scales and phenotypic information by trained fishers^15,33^. Scales were stored in envelopes at room temperature and used to determine individual life-history characteristics including the number of years spent in the freshwater environment prior to smoltification (river age), number of years spent in the marine environment prior to maturation (sea age) and possible previous spawning events, following international guidelines (ICES 2011). The Teno river Atlantic salmon have diverse life history strategies ^15,35^. They can spend from two to eight years in freshwater before smoltifying, from one to five years at sea before maturing and have up to five breeding attempts. Overall, a total of 120 combinations of river age, sea age at maturity and repeat spawning strategies have been described^15^. Age at maturity has been declining in Teno salmon over the last 40 years, with proportionally fewer late maturing salmon returning over years. Age at maturity also differs largely among populations displaying genomic signatures of local adaptation^15,33^.

We randomly selected scales from individuals caught by rod between 1972 and 2014 during the later part of the fishing season, from July 20 to August 31. Most of the Teno salmon are expected to have reached their home river by late July ^36^. Samples came from two different locations, the middle reaches of the Tenojoki mainstem (hereafter Tenojoki) and a headwater region Inarijoki (Supplementary Figure 5). These sections of the river host weakly differentiated salmon populations with contrasting sea-age structure at maturity^31,33^. Individuals from the Tenojoki spend, on average, more time at sea before maturing than individuals from the Inarijoki population ^15,30^. Seventy additional females were selected in Inarijoki over the study period, by following the same sampling scheme, to increase the sampling size in analyses with sex-specific estimates. Scale or fin samples were also collected from juvenile salmon from the Tenojoki (N=143, 2–3 years old) and Inarijoki (N=108, 1–3 years old) populations caught by electrofishing in the 2012 and 2016, respectively. They were used as the baseline for population assignment of adults and to determine potential sex-specific *vgll3* allele frequency differences at the juvenile stage.

### Genotyping

DNA extraction from scales, sex determination and genotyping were performed following^37^. In total, 2482 individuals were genotyped at 191 SNPs, including the SNP the most highly associated with the age at maturity, *vgll3TOP* (vestigial-like family member 3 gene also called *vgll3*,^10^) and outlier and baseline SNP modules^37^. The outlier module consisted of 53 SNPs highly differentiated between the Inarijoki and Tenojoki populations, thus allowing a more powerful assignment of population of origin, between these two closely related populations ^i.e. see 30,37^.The baseline module included 136 putatively neutral markers in low linkage disequilibrium, distributed over the whole genome proportionally to chromosome length, previously filtered to have minor allele frequency >0.05 and heterozygosity >0.2^37^. These SNPs were used to estimate the level of differentiation among populations of the Teno River (Weir and Cockerham’s F*_ST_*) and genetic drift. Mean genotyping success was on average 0.80 per locus and individual.

### Population assignment

The 53 outlier loci were used to determine the optimum number of genetic clusters and assign the population of origin of adults using the software STRUCTURE. First, an admixture model with correlated allele frequencies^38^ was run on adult and juvenile data for 80,000 MCMC iterations, including a burn-in length of 50,000. The model was replicated six times for each cluster value K, varying from one to four. The optimal number of clusters was thereafter estimated using the Δ K method described in Evanno, Regnaut, and Goudet (2005) using STRUCTURE HARVESTER ^40^. This allowed us to determine whether juveniles were correctly assigned to their sampling locations and could thus be used as a baseline for adult assignment. Then another admixture model with correlated allele frequencies was replicated six times on adult data using juvenile data as a baseline, with prior migration set to 0. The *fullsearch* algorithm from the CLUMP software (Jakobsson & Rosenberg 2007) was used to account for across replicate variability in membership coefficients. Finally, the cluster of each adult was assigned by using the optimum K and membership probability superior or equal to 0.8. The differentiation between populations was tested by calculating the likelihood ratio G-statistic ^41^ and comparing it with the G-statistic distribution obtained by permuting 1,000 times individuals between populations.

The most likely number of clusters determined with the Δ K method was two when juveniles and adults data were combined (Supplementary Figure 6 Supplementary Figure 6). Juveniles were assigned accordingly to their sampling location in more than 96% of cases (Supplementary Figure 7). Using juvenile data as a baseline, 90% of adults were classified to one of the two clusters with probabilities equal or higher than 0.8. Individuals sampled in Tenojoki were assigned to the Inarijoki population in 25% of the cases whereas only 2% of the individuals caught in Inarijoki were assigned to the Tenojoki population. In total, 1330 and 911 individuals clustered in the Tenojoki and Inarijoki populations, respectively (Supplementary Figure 8). The two populations were significantly genetically differentiated (*F_ST_* = 0.013, *G* = 201.55, P < 0.01) and had contrasted age structures (Supplementary Figure 9).

### Statistical analyses

#### Temporal variation in age at maturity and proportion of females

Non-linear temporal variation in age at maturity was estimated separately for each population using generalized additive models, with the Gaussian family as the residual distribution. Year of hatching was included as an independent variable inside a cubic regression spline for each sex. The study included spawning individuals caught over a 43 year period (1972 to 2014). Hatch years were calculated based on the specific life-history strategy of each individual and spanned the period from 1971 to 2006 in Tenojoki and 1971 to 2007 in Inarijoki. Sex was also included as an explanatory variable.

The amount of smoothing was determined in each case using the maximum likelihood method. Automatic smoothness selection was performed by adding a shrinkage term. The significance of independent variables was assessed using F-tests and an alpha risk of 0.05. All statistical tests included in this manuscript were two-tailed. The additive models were run with the *R* package *mgcv* ^42,43^.

#### Effect size of *vgll3* on age at maturity

To estimate the genetic effect of *vgll3*, age at maturity was also regressed using a multinomial model separately for each population. In Tenojoki, two individuals having matured after five years at sea were considered having matured after four years to avoid the estimate of additional model parameters without data support. The sex, year of capture and *vgll3* genotype can all influence age at maturity and were included in models as a three-way interaction. Multinomial models in this study were performed using the *R* package nnet ^44^. Model selection was performed using backward selection with F-tests and by calculating the AICc of all possible models. The effect of year on the probability to mature was calculated with the Effect package ^45^ which averages the effect size across sexes and genotypes. The mean age at maturity per sex and genotype was calculated from model predicted values. First, predicted age was obtained for each year, sex and age at maturity combination by multiplying the probabilities of having matured after one, two, three or four years at sea by the corresponding sea age at maturity and taking the sum. Second, the age at maturity was calculated for each sex and genotype by averaging over years. This process was replicated 1000 times by randomly sampling with replacement and fitting a new model. A 95% bootstrap confidence interval was then determined by taking values of the 2.5 and 97.5 percentiles. The *vgll3* alleles were called *L* and *E* to indicate their association with late and early maturation, respectively ^10^. Dominance for each sex and population was estimated from the mean age at maturity (μ) following 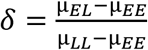. The *L* allele is recessive if *δ* = 0, additive if *δ* = 0.5 and dominant if *δ* = 1.

To determine how much of the observed changes in age at maturity over time could be attributed to changes in genotypes and year of capture, a new dataset with the spawning year held constant at 1975 was created for Tenojoki. The previous multinomial model was used to predict new maturation probabilities from which model predicted age at maturity were calculated for each individual, as above. Temporal changes in age at maturity attributed to genotypes were determined by fitting a generalized additive model using the Gaussian family and including the individual hatch year in a cubic regression spline and the sex as independent variable. Changes in age at maturity attributed to the year of capture corresponded to the difference between individual predicted age at maturity calculated from the original dataset and the one with the year fixed. Another Gaussian generalized additive model was also performed on those differences, by including the hatch year in a cubic regression spline. Automatic smoothness selection was performed by adding a shrinkage term.

#### Change in allele and genotype frequencies

Temporal variation in allele frequencies was determined for each population and locus using generalized linear models (*glm*), with the quasibinomial family to account for overdispersion. Sex-dependent *vgll3* genetic effect on the age at maturity ^10^ may create sex-specific selection at sea, leading to differences in *vgll3* allele frequency between male and female spawners from the same generation. The sex variable can capture this potential intra-generation variation in allele frequency. Hence, sex and year of hatching were included as independent variables in the *glm*. To keep the potential effect of sex-specific selection on the *vgll3* allele frequency temporal change, the model was also run without including sex as a covariate. The significance of variables was assessed with F-tests.

In order to determine whether *vgll3* allele frequencies varied across time more than under the neutral expectation, model predicted temporal changes in allele frequencies were compared among loci with individual genotyping success higher than 0.7 (144 and 135 loci for the Tenojoki and Inarijoki populations, respectively). This threshold was chosen as a trade-off between increasing the quality and amount of data per locus (average genotyping success superior to 0.90 in those subsets) and keeping a large number of loci for the comparison (~25–30% of loci were excluded). The amount of genetic drift, and thus random temporal allele frequency change, is dependent on the initial allele frequency of each locus ^46^. The comparison of temporal changes between *vgll3* and other putatively neutral loci was thus corrected for initial allele frequency by calculating the expected amount of drift at *vgll3* under a Wright-Fisher model ^47,48^. The distribution of allele frequency *x*(*t*) after t generations can be approximated using a normal distribution ^46^:

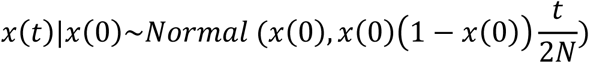

with *x*(0) being the initial allele frequency and *N* the effective population size. The ratio 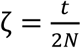 was estimated with a Bayesian model using a uniform prior distribution ranging from 0 to 1. The binomial fitted allele frequencies at birth for years 1971 (*x*(0)) and 2006 (*x*(*t*)) in Tenojoki and 1971 (*x*(0)) and 2007 (*x*(*t*)) in Inarijoki were used as data for all loci except *vgll3*. Two MCMC chains were run for 100,000 iterations with thinning interval 10, and a burn-in length of 100,000. Convergence was assessed using the Gelman and Rubin’s convergence diagnostic^49^ and a potential scale reduction factor (*psrf*) threshold of 1.1. The probability to observe the *vgll3* allele frequency change (*Δ_vgll3_ =|x*(*t*)*_vgll3_ − x*(*0*)*_vgll3_|*) under drift alone was calculated at each of the 20,000 saved iterations (*i*) to account for uncertainty in ζ estimation as follows:

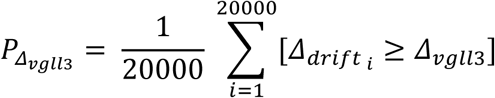

with

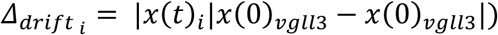

and

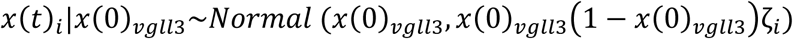

To account further for uncertainty in *vgll3* allele frequency change, *Δ_vgll3_* was re-estimated at each iteration *i* by running the quasi-binomial model with a new dataset, sampled for each year from the original dataset with replacement. The distribution of changes expected under drift alone was also calculated in the same manner as for *vgll3*, for initial allele frequencies *x*(0) varying from 0.5 to 1.

To determine whether potential differences in *vgll3* allele frequencies between adult males and females are likely to arise during the sea migration, juvenile allele frequencies were analyzed using a separate *glm* with the binomial family for each population. Sex was introduced as an independent variable. A backward model selection was performed using Likelihood-Ratio Tests (LRT) and an alpha risk of 0.05. Confidence intervals were calculated with the lsmean package ^50^ by taking the years 2006 and 2007 as reference for the Tenojoki and Inarijoki adults, respectively.

To further describe temporal changes in *vgll3* genotypes, a multinomial model was used for each population. The year of hatching and the sex were introduced as independent variables in a two way interaction. A backward model selection was performed using LRT and an alpha risk of 0.05.

Allele frequencies may differ between sexes due to sex-specific selection between homozygotes but also because of the effect of sex-specific dominance, even when selection is sex-independent. To determine how dominance can contribute to differences in allele frequencies between sexes in the latter case, the expected sign and magnitude of allele frequency differences were determined for different selection strengths, by using the dominance patterns calculated previously for the Inarijoki and Tenojoki populations. Considering a gene with 2 alleles *A* and *B* with respective frequencies *p* and *q*, the allele frequency after a selection event corresponds to:

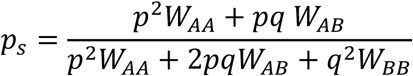

with *W_AA_, W_AB_* and *W_BB_* the relative fitness of each respective genotype:

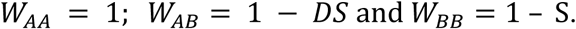

where S is the selection coefficient common to each sex, varying from 0 to 0.90 by 0.15 intervals. *D* is the dominance coefficient. *P_s_* was calculated for each sex and population using the corresponding dominance coefficients previously calculated from phenotypes (*δ*) and an initial *p* varying from 0 to 1. The expected difference in allele frequency in Supplementary Figure 2 corresponds to *p_s_*(female) - *p_s_*(male), calculated for each combination of S and *p*.

#### Estimation of selection coefficients

A Bayesian model was used to estimate selection coefficients by accounting for drift induced by a limited number of spawners, in a similar way to Wright-Fisher models ^e.g. 51,52^ First, the linkage disequilibrium method ^53^ implemented in the software NeEstimator 2.01 ^54^ was applied on samples grouped by cohort year to estimate the parental effective number of breeders (*N*b). This approach was favored over the standard temporal method potentially generating biased effective size estimates when used with temporally close samples from species with overlapping generations ^55^ and only providing information about the harmonic mean of effective sizes. In order to use the linkage disequilibrium *Nb* values and associated 95% parametric confidence intervals in the Bayesian models, parameters of log-normal distributions with similar percentiles were assessed using the R package *rriskDistributions* ^56^. Weights of 7, 2 and 1 were respectively assigned to the 2.5, 5.0 and 97.5 percentiles to increase the approximation precision for lower bounds and medians. The negative or infinite values were replaced by 5000 or 10 000 for the median and 95% confidence interval upper bound, respectively. These are realistic maximum breeder numbers in the populations and represent a conservative approach. If the lower bound also displayed infinite values, the corresponding distribution had a median of 9 000 and lower and upper bounds of respectively 8 000 and 10 000.

The selection coefficient represents “the reduction in relative fitness, and therefore genetic contribution to future generations, of one genotype compared to another”^57^. Selection coefficients were estimated using 32 and 33 different spawning years, with corresponding hatch years, for Tenojoki and Inarijoki, respectively. Considering a SNP with alleles A1 and A2 and 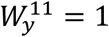, 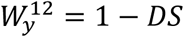 and 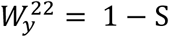 being the relative fitness of each genotype. S corresponds to the selection coefficient, following a uniform prior distribution ranging from −1 to 1. *D* denotes the dominance coefficient, following a uniform prior distribution ranging from 0 to 1. The observed number of each genotype *g* in spawners of sex *s* in year *y* (*O_g,s,y_*) followed a Dirichlet Multinomial (DM) distribution:

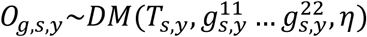

where *η* is the variation parameter following a uniform distribution ranging from 1 to 2500 and *T_s,y_* the total number of spawners per sex and year. The spawners genotype frequency for each sex 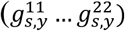 varied over years according to a hierarchical model, 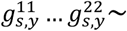 dirichlet 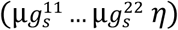 and 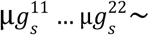 dirichlet (1,1,1). The observed number of allele A1 (*n_y_*) in individuals born in year y follow a binomial distribution:

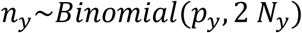

with *N_y_* being the total number of individuals per year and *p_y_* the population allele frequency. The expected allele frequency in the cohort y depends on genotype frequency in spawners the year before as follows:

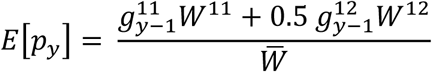

with 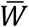 being the population mean fitness 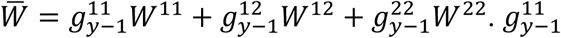, 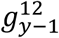 and 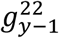 are the genotypes of spawners averaged across sexes, as each sex contributes equally to the next generation despite a potential biased sex-ratio. Genetic drift should be taken into account to estimate p_y_ from the genotype frequencies of the previous year’s spawners. In populations with random mating, it corresponds to drawing randomly p_y_ from a binomial distribution ^46–48^ with as parameters the expected allele frequency E[py] and twice the effective number of spawners, previously estimated with the linkage disequilibrium method (2 *Nb_y_*). Consequently, the expected variance of the allele frequency *p_y_* subject to drift is after one generation 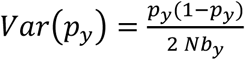. For computing time and convergence reasons, a beta distribution with equal mean and variance was used instead:

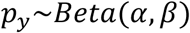

with

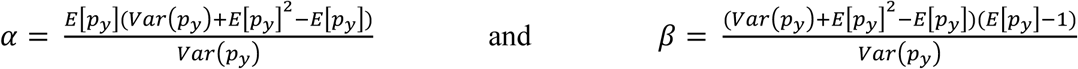

Priors used in this model were chosen to be as uninformative as possible. For the *vgll3* locus, the *L* allele was chosen as reference. The “pMCMC” were calculated from the two chains as following: 2 * min(p < 0; 1 - p < 0), p<0 being the proportion of values below zero.

Posterior distributions were approximated using Monte Carlo Markov Chain (MCMC) methods with the Just Another Gibbs Sampler software (JAGS^58^) run in the R environment^43^. Two MCMC chains were run for 4.5 million iterations, including a burnin length of 3.5 million. Only one iteration out of 100 was kept to reduce the memory size used. Gelman and Rubin’s convergence diagnostic ^49^ was used to assess convergence. Models were run longer if the potential scale reduction factor (*prsf*) was initially superior to 1.10. Finally, all models had potential scale reduction factor inferior or equal to 1.10 for all parameters, except for up to 2 *Nb_y_* parameters in 10 models for Inarijoki, having larger *psrf* (inferior to 1.30).

#### Data and code availability

The datasets used during the current study will be uploaded to a public data repository upon acceptance.

#### Code availability

The custom codes used during the current study are available from the corresponding author on reasonable request.

## Acknowledgements

We thank numerous fishers who participated in the collection of scales and phenotypic information, Eero Niemelä for starting the program and looking after contacts with fishers over the 40 year study period, Jorma Kuusela for organizing the samples collection from the archive and several scale readers, especially Jari Haantie. This project received funding from the European Research Council (ERC) under the European Union’s Horizon 2020 research and innovation programme (grant agreement No 742312) as well as from the Academy of Finland (projects No. 284941, 286334, 307593, 302873 and 318939)

## Author contributions

J.E and P.O. coordinated the collection of samples; C.R.P., Y.C., T.A. and J.E. designed the study; Y.C. analyzed the data; Y.C., C.P. and T.A. wrote the manuscript and all authors contributed to its revision.

## Competing interests

The authors declare no competing financial interests.

Correspondence and requests for materials should be addressed to C.R.P.

